# A synthetic membrane shaper for controlled liposome deformation

**DOI:** 10.1101/2021.12.22.473854

**Authors:** Nicola De Franceschi, Weria Pezeshkian, Alessio Fragasso, Bart M.H. Bruininks, Sean Tsai, Siewert J. Marrink, Cees Dekker

## Abstract

Shape defines the structure and function of cellular membranes. In cell division, the cell membrane deforms into a ‘dumbbell’ shape, while organelles such as the autophagosome exhibit ‘stomatocyte’ shapes. Bottom-up *in vitro* reconstitution of protein machineries that stabilize or resolve the membrane necks in such deformed liposome structures is of considerable interest to characterize their function. Here we develop a DNA-nanotechnology-based approach that we call Synthetic Membrane Shaper (SMS), where cholesterol-linked DNA structures attach to the liposome membrane to reproducibly generate high yields of stomatocytes and dumbbells. *In silico* simulations confirm the shape-stabilizing role of the SMS. We show that the SMS is fully compatible with protein reconstitution by assembling bacterial divisome proteins (DynaminA, FtsZ:ZipA) at the catenoidal neck of these membrane structures. The SMS approach provides a general tool for studying protein binding to complex membrane geometries that will greatly benefit synthetic cell research.

## Introduction

Biological membranes constitute the chassis of cells in all living organisms, providing structural support, compartmentalization, and a platform for organizing biochemical reactions. A fascinating property of membranes is their ability to adopt a variety of shapes, a feature that is exemplified in the rich repertoire of morphologies observed in intracellular organelles.^1^ The Golgi apparatus, for instance, is composed of a stack of cisternae, a membrane arrangement that optimizes the stepwise post-translational modification of proteins that are processed in the organelle. Two shapes constitute particularly important membrane geometries in cells (Figure 1A, S1A): The dumbbell shape mimics the geometry of a dividing cell, while the stomatocyte shape recapitulates the membrane topology found in several intracellular organelles including the nuclear envelope and the open autophagosome. A common feature that is shared by both dumbbells and stomatocytes is a neck, a double-membrane pore with the geometrical shape of a catenoid that features both positive and negative membrane curvature (Figure 1A). Membrane deformations result from the combined effects of the spontaneous and induced local curvatures^2^. The molecular origin of spontaneous curvature can be explained by a number of mechanisms. It can, for example, result from asymmetries in lipid composition in the bilayer, from bulky groups that insert into one leaflet of the membrane (‘wedges’), from oligomerization of membrane proteins that build up complexes with an intrinsic curvature (‘scaffolding’), and from molecular crowding due to entropic repulsion of soluble protein domains arising from confinement in a crowded environment. From previous studies it appears that scaffolding and wedging are the most effective ways to induce curvature, with crowding having a more modest effect^3,4^.

**Figure 1:**
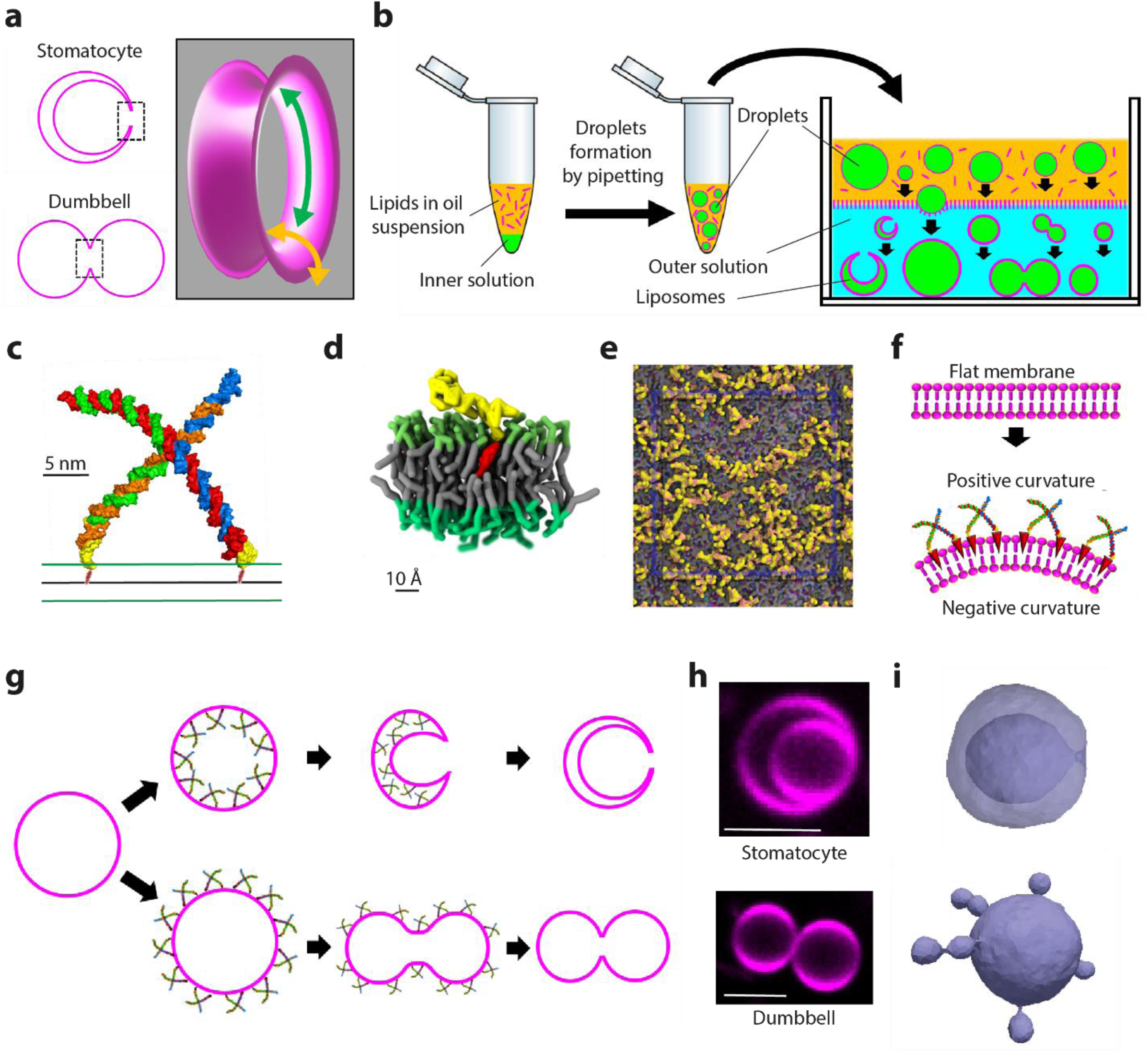
Membrane shaping by the SMS approach. **(a)** Schematics showing the toroidal geometry of a double-membrane pore (right) present in both stomatocyte (top) and dumbbell shapes (bottom). Negative and positive membrane curvatures are indicated by the green and orange arrows, respectively. **(b)** Schematics depicting liposome production with the SMS approach. Water-in-oil droplets are produced by pipetting and then deposited in the observation chamber, were they sink by gravity. The bilayer is created when the droplets cross the interface between oil and aqueous (outer) solution. An osmotic pressure difference between inner and outer solutions induces shape deformations in the newly formed liposomes. Inner solution is denoted in green, oil phase in orange, lipids in magenta, and outer solution in cyan. **(c)** Structural model of a single nanostar formed by four ssDNA oligos (red, blue, orange, green) and hybridized to the chol-oligos (yellow). The position of the cholesterol moiety (red) embedded within the membrane bilayer (green and black lines) is schematically indicated. **(d)** Representative snapshot of an all-atom simulation of a chol-oligo inserted into the membrane (color-coding as in panel C). **(e)** Top view snapshot of an all-atom simulation of chol-oligos inserted onto a POPC/Chol membrane. **(f)** Schematic illustrating curvature generation by the combination of nanostars and chol-oligo. **(g)** Shape transformations resulting from assembling the nanostars on either the inner or the outer side of liposomes. **(h)** Example of a confocal image of stomatocyte and dumbbell shaped liposomes generated by the SMS. Scale bars: 5µm. **(i)** Stomatocyte and dumbbell obtained by DTS simulations.

An elegant body of knowledge has been developed with theoretical calculations describing membrane shape transformation of spherical liposomes into a wide variety of shapes as a result of varying the membrane spontaneous curvature, surface-to-volume ratio, and bulk parameters such as temperature^5^. Experimentally, membrane deformation has been extensively studied in osmotically deflated liposomes^2^. Dumbbell-shaped vesicles have, for example, been obtained by asymmetric absorption of sugars^6^, large DNA molecules^7^, dextran^8^ and proteins^9^, as well as induced by Min protein activity^10^ or active particles^11^. Stomatocyte-like shapes have, however, rarely been reported in literature – merely as short-lived intermediates in “vesicles-in-vesicles” membrane systems^12^. The ability to reproduce these shapes *in vitro* for quantitative characterization of the assembly and function of protein complexes on their neck regions would be beneficial for bottom-up synthetic biology. In dumbbells, for example, the neck represents the assembly site for the division machinery, while in stomatocytes, the neck is the assembly site of the nuclear pore complex at the nuclear envelope as well as the membrane topology where the ESCRT-III complex binds.

Due to the unavailability of good model systems, only sporadic examples^12^ of such protein reconstitutions are available^13^. A common drawback of experimental approaches is the requirement for specific lipid and buffer compositions that restrict their applicability to study biological processes. Furthermore, comparison between different membrane geometries is hardly possible since each system is designed to obtain one specific shape, and poor control and low yields limit quantitative characterization of complex membrane shapes.

Here, we establish a novel approach that we name the “Synthetic Membrane Shaper” (SMS) to shape liposomes in a highly controlled way. The SMS is a method in which small DNA nanostructures adsorb onto the surface of a liposome that is being formed in a hyperosmotic environment. This induces and stabilizes large-scale membrane deformations, generating both dumbbell and stomatocyte shapes. The SMS works within a broad range of membrane and buffer compositions and is widely applicable, as we show by assembling various proteins on high-curvature regions such as necks. We use *in silico* dynamically triangulated surfaces (DTS) simulations^14,15^ to verify the vesicle shape transformations and characterize the assembly of proteins to regions of complex membrane curvature.

## Results

### Establishing a synthetic membrane shaper to morph liposomes into dumbbells or stomatocytes

Liposomes were generated in a gentle method where water-in-oil droplets slowly crossed an oil/water interface by gravity (Figure 1B). This method was chosen to overcome harsh procedures in many common production and handling techniques for liposomes that, due to centrifugation and pipetting, can cause loss of delicate membrane structures. Droplets were prepared by pipetting the inner aqueous solution into a lipid-in-oil dispersion, whereby water-in-oil droplets with a lipid monolayer were formed. Subsequently, these droplets were transferred by gravity through an oil/water interface to acquire a second bilayer leaflet and thus form liposomes in a hyperosmotic outer solution in an observation chamber, where liposomes settled at the bottom where they were imaged with fluorescence microscopy. With this liposome-production technique, it is very easy to include macromolecules such as proteins or DNA in both the inner and outer solutions, while preserving fragile membrane structures. Membrane shape transformation are driven by the osmotic difference between the inner and outer solutions (20-40 mOsm in our study). This resulted in membrane deformations that however were not greatly controlled, with both the total yield and the relative occurrence of different shapes varying greatly across preparations. A variety of morphologies was observed, including dumbbell and stomatocyte shapes, sometimes even coexisting in the same preparation (Figure S1B).

In order to increase the yield and reproducibility of structures, we developed an approach to drive vesicles into a defined shape. We used DNA nanostars^16^, which are 96.5kDa tetrameric cross-shape DNA assemblies (Figure 1C) that at two positions have a binding site to a short complementary oligonucleotide (chol-oligo) that is functionalized with a cholesterol moiety at its 3’ end. Insertion of the chol-oligo into membranes is expected to generate membrane curvature by the wedge effect, while the nanostars are expected to induce a further bilayer asymmetry by molecular crowding. Molecular Dynamics (MD) simulations of insertion of the chol-oligo into membranes indeed indicated that such DNA strands, that are covalently attached to cholesterol that is dipping into the hydrophobic core of the upper leaflet of the bilayer (Figure 1D, 1E), displace the lipid headgroups laterally. This wedge effect effectively increases the area of the upper layer and addition of the nanostars/chol-oligo complexes can thus be used as a tool to induce membrane curvature by bending the membrane away from the side where the nanostars were bound (Figure 1F).

This combination of the action of the DNA nanostars/chol-oligo complex with the gentle liposome production method constitutes an approach that we name Synthetic Membrane Shaper (SMS). As we detail below, this SMS approach was found to enable the production of two kinds of membrane shapes: stomatocyte-shaped liposomes when the nanostars were bound to the inner leaflet, and dumbbells for binding of the nanostars to the outer leaflet (Figure 1G, H). DTS simulations showed that merely the combination of a mild osmotic pressure difference and membrane curvature (here induced by the nanostars/chol-oligo) can result in such membrane deformations. Stomatocyte-like structures were generated upon applying negative global membrane curvature on liposomes with a low constant volume, while dumbbell shapes were similarly generated by a positive membrane curvature (Figure 1I, S2, S3; movies 1 and 2).

### Characterization of stomatocytes and dumbbells produced with the SMS

The presence of nanostars in the inner aqueous solution of the liposomes strongly increased the yield of stomatocytes, i.e., from 7% without to 78% with nanostars (Figure 2A). Nanostars were found to be distributed homogenously throughout the membrane surface without clustering at the microscale (Figure 2B). MD simulations indicated that the chol-oligos also did not form stable clusters at the nanoscale (Figure S4). The SMS approach to induce shape deformations appeared to work efficiently on liposomes with a size in the lower µm range (Figure 2C), while larger vesicles largely remained spherical. We also separately investigated the contributions of the chol-oligos and nanostars in mediating membrane deformation (Figure S5A, S5B).

**Figure 2:**
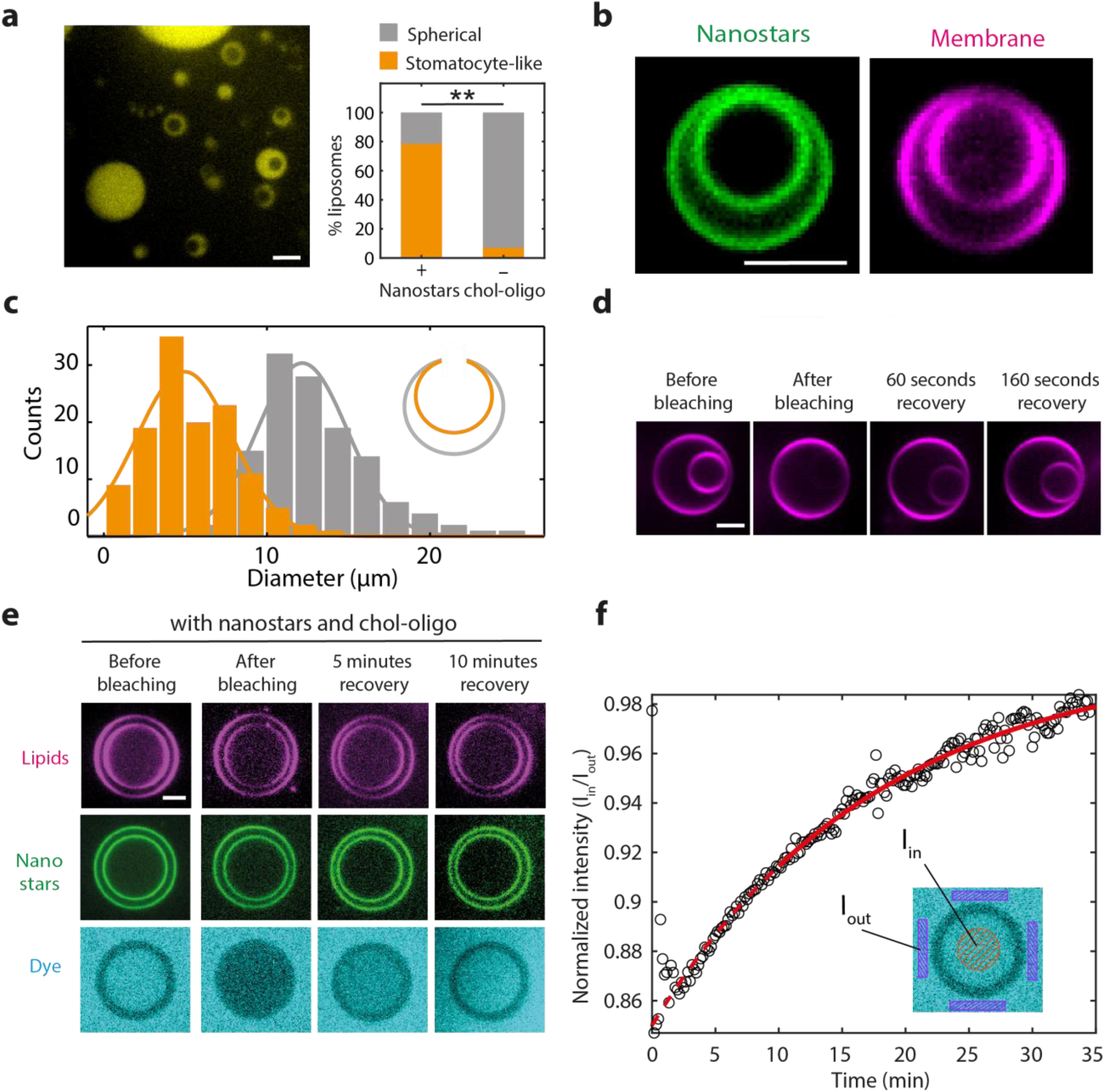
Characterization of stomatocytes produced with SMS. **(a)** Example of a field of view in a SMS stomatocyte preparation. A soluble dye (visible in yellow) was encapsulated in the lumen of the stomatocytes. The plot (right) shows the quantification of the number stomatocytes obtained with or without nanostars/chol-oligo (N=511 liposomes from 3 independent preparations). **(b)** Representative example of a stomatocyte, imaged both in the lipid (magenta) and nanostars (green) fluorescence. Nanostars were observed to be uniformly distributed over the membrane. **(c)** Distribution of the diameter of the inner and outer membrane of stomatocytes. Outer diameter was 13.1 ± 3.7 µm (average±SD); inner diameter was 5.4 ± 2.7 µm (average±SD). N=125 stomatocytes from 5 independent preparations. **(d)** FRAP experiment on fluorescent lipids. The inner membrane is bleached and fluorescent recovery over time is visualized. **(e)** FRAP experiment of soluble alexa-647 dye contained in the lumen of stomatocyte obtained by the SMS approach in the presence of nanostars/chol-oligo. Stomatocytes were produced in the presence of alexa-647. The dye present in the inner compartment is photobleached and its recovery through the toroidal pore is visualized over time. **(f)** Plot showing the normalized intensity of the dye in function of time. The inset depicts the regions where I_in_ (fluorescence intensity in the inner compartment, in red) and I_out_ (fluorescence intensity in the outer buffer, in purple) were measured. Solid line denotes a fit of Eq. 7 (Supplementary note 1). All scale bars: 5µm.

By optical imaging alone it cannot be determined whether the neck connecting the inner and outer membrane in stomatocytes is open. Indeed, while in few instances an elongated neck could be imaged (Movie 3), the size of the neck was below the diffraction limit in the vast majority of stomatocytes. We therefore performed a series of FRAP (Fluorescence Recovery After Photobleaching) experiments to validate the presence of an open neck. After photobleaching the fluorescent lipids of the inner vesicle, we visualized the fluorescence recovery and equilibration of membrane lipids between inner and outer vesicle (Figure 2D). Furthermore, we generated stomatocytes in the presence of a soluble dye in the outer solution. As expected, the dye was observed in both the inner vesicle and the exterior of the stomatocytes right after production. We then diluted the external solution with buffer that did not contain dye, thereby testing whether the dye would exit the inner vesicle through an open pore, or would be retained in case the pore was closed. In this way, we estimated that the majority of stomatocytes (55%) clearly exhibited an open neck (Figure S5C).

FRAP experiments also allowed to estimate the neck size. Figure 2E shows a characteristic example of a FRAP recovery in a stomatocyte, where first the inner vesicle was photobleached, followed by recovery of the small soluble dye (Alexa488) within tens of minutes. By fitting a diffusion model (as described in Ref.^17^, Supplementary Note 1 and Figure S6) to the data (Figure 2F), we could estimate an open pore diameter of 134 ± 103 nm (N=9, error is SD). This is remarkably smaller compared to the control experiment on stomatocytes that were formed in absence of nanostars and chol-oligo, where recovery occurred within tens of seconds, corresponding to a pore size of 1.3±1.2 μm (N=8, error SD; see Figure S7). This indicates that the chol-oligo and nanostars indeed generate membrane curvature, resulting in more constricted necks.

Liposome SMS preparations where nanostars and chol-oligo were added on the *outside* were found to be strongly enriched in dumbbells and chains of dumbbells (Figure 3A,B). The diameter of the lobes in these dumbbells were again limited to the low µm range (Figure 3C), as also observed for the stomatocytes. In the majority (81%) of dumbbells observed, FRAP analysis indicated that lipids did flow across the neck, confirming that the lipid membranes of adjacent lobes were connected (Figure 3D and Movie 4). Accordingly, when performing FRAP experiments on chains of dumbbells, we observed fluorescent-lipid recovery that proceeded sequentially from each lobe to the next (Figure 3E). Upon encapsulation of soluble dye within the lumen of these dumbbells and photobleaching of one lobe, we observed dye recovery (Figure 3F), indicating that the lumens of adjacent lobes are in mutual communication via an open neck. Figure 3F shows fitting of the FRAP recovery data to a diffusion model that describes the molecular flow in a dumbbell two-vesicle system (see Supplementary Note 2 and Figure S8), from which we estimate an open pore diameter of 26 ± 23 nm (N=7, error is SD). The data indicate that the large majority of the structures that we obtained were indeed true dumbbells rather than individual liposomes that were adhering to each other *post hoc*.

**Figure 3:**
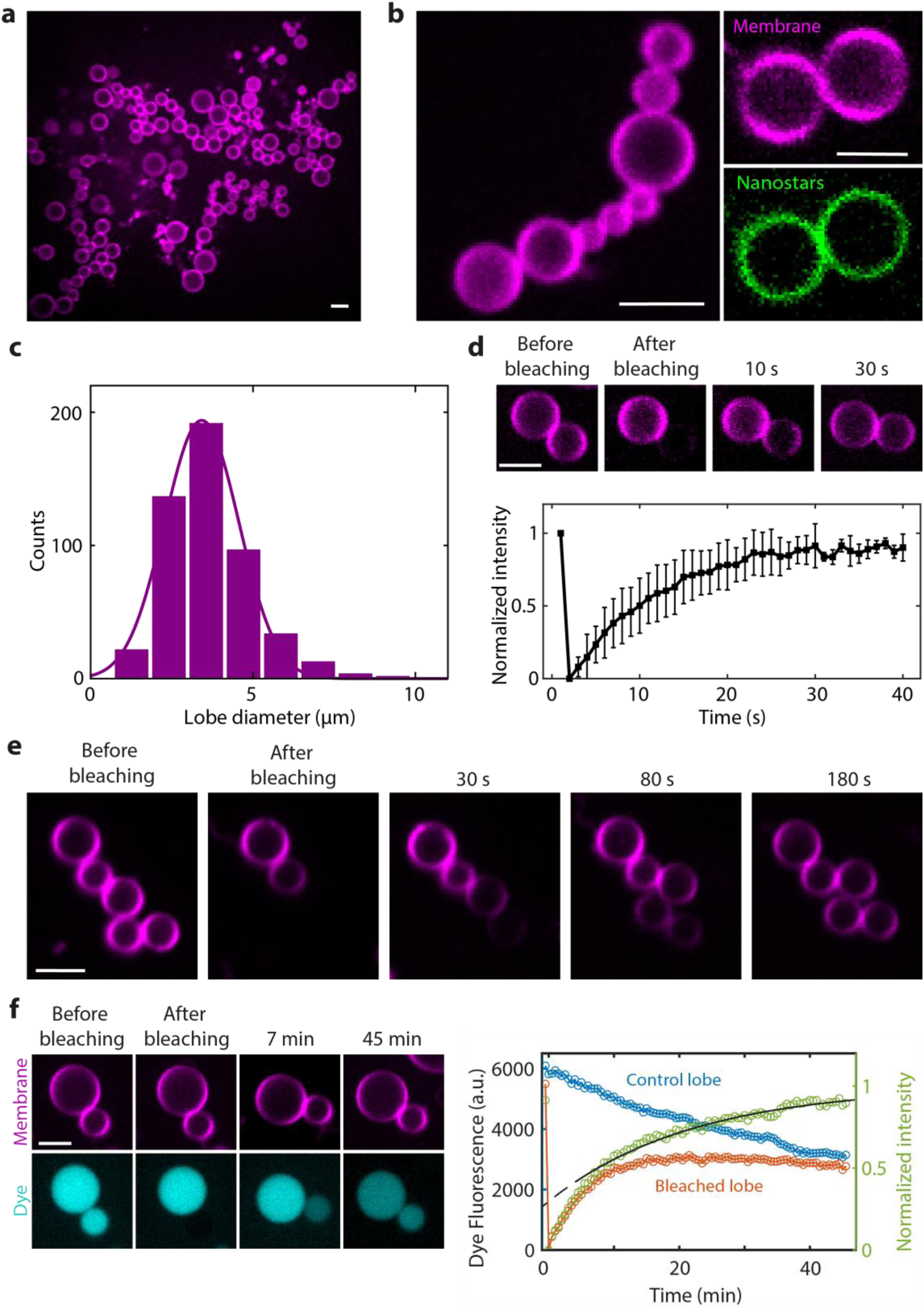
Characterization of dumbbells produced with SMS. **(a)** Large field of view in a SMS dumbbell preparation. **(b)** Example of a chain of dumbbells (left) and a dumbbell (right). The nanostars (green) are observed to be homogeneously distributed throughout the membrane. **(c)** Quantification of dumbbell lobe diameter. N=501 lobes from 6 independent preparations. **(d)** Recovery of fluorescent lipids upon FRAP photo-bleaching of one of the two lobes of dumbbells. The plot shows the average recovery time of the fraction of dumbbells (81%) that exhibited full recovery. N=31 dumbbells from 6 independent preparations. **(e)** FRAP recovery of fluorescent lipids upon photobleaching of part of a chain of dumbbells. **(f)** FRAP experiment showing flowing of soluble dye between lobes of a dumbbell. The right plot shows the fluorescent intensity versus time. Solid line denotes a fit of Eq. 16 (Supplementary note 2). All scale bars: 5µm.

### Dynamin A and FtsZ:ZipA proteins assemble at the necks of stomatocytes and dumbbells

Subsequently, we used the SMS approach to characterize protein assembly at necks of dumbbells and stomatocytes. We first tested the assembly of Dynamin A (DynA). This bacterial member of the dynamin superfamily localizes to the neck of dividing cells where it has been proposed to mediate the final step of membrane scission^18^. However, thus far DynA could never be reconstituted and imaged *in vitro* on membrane necks. We reconstituted DynA in both stomatocytes and dumbbells obtained by SMS. Note that in both cases, the protein and the nanostars are bound to opposite sides of the membrane (see schematics in Figure 4A and 4D) and hence they will not mutually interact.

**Figure 4:**
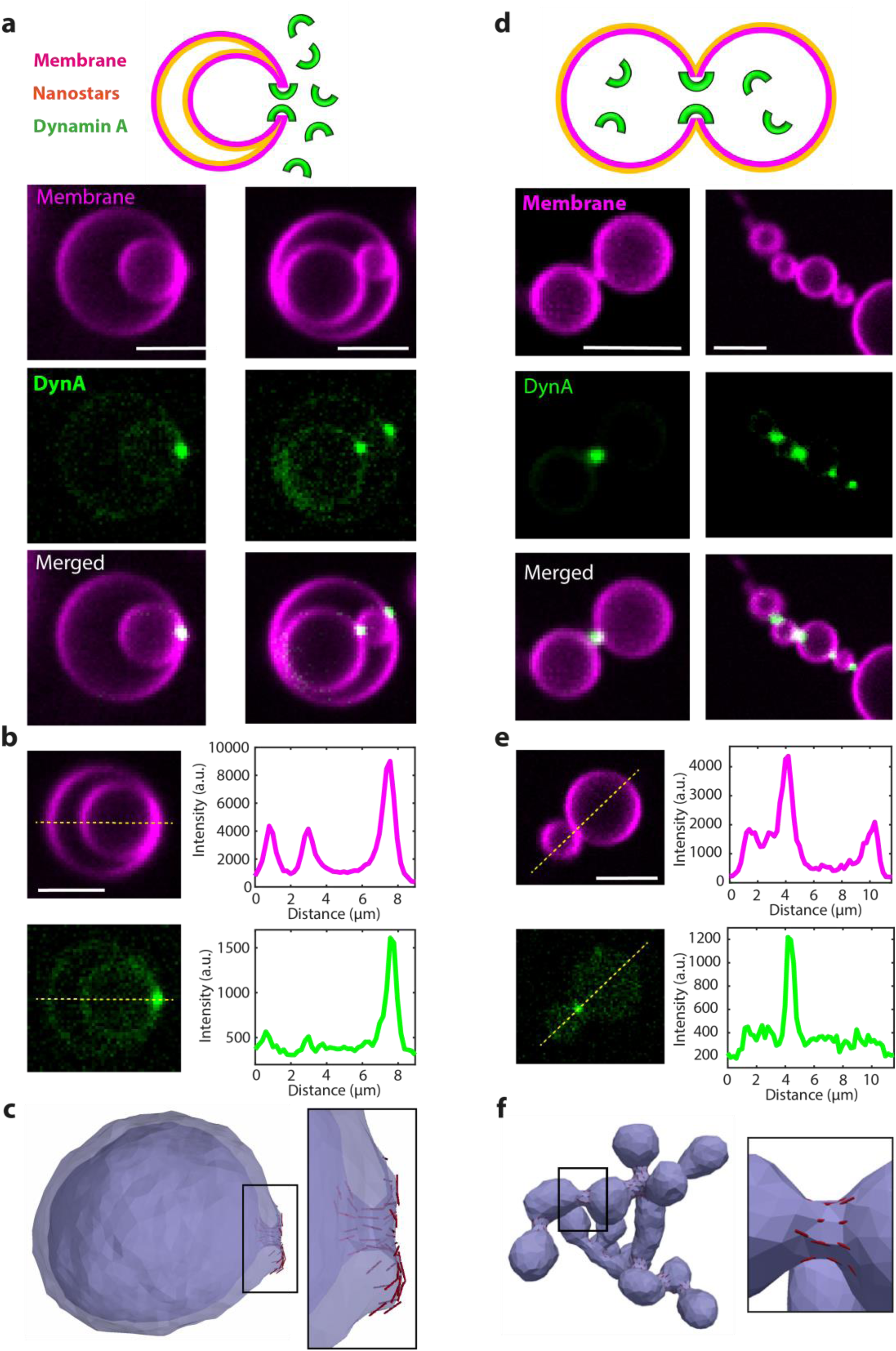
Assembly of Dynamin A on stomatocyte- and dumbbell-shaped liposomes. **(a)** Assembly of a Dynamin A on stomatocytes generated by SMS. Schematic on the top row clarifies the topology of the protein and the nanostars with respect to the membrane. Lower panels show representative images from 3 independent preparations. **(b)** Line scan analysis of Dynamin A enrichment at necks of stomatocytes. **(c)** DTS simulation of a vesicle under osmotic pressure and negative global curvature and 5% protein bound on the outside. Each red line represents one protein and its orientation on the membrane. Proteins induce positive membrane curvature. **(d)** Assembly of Dynamin A inside dumbbells generated by SMS. Schematic on the top row clarifies the topology of the protein and the nanostars with respect to the membrane. Lower panels show representative images from 3 independent preparations. **(e)** Line scan analysis of Dynamin A enrichment at necks of dumbbells. **(f)** DTS simulation of a vesicle under osmotic pressure and positive global curvature and 5% protein coverage. The proteins, which are bound to the membrane from the inside, induce positive membrane curvature. All scale bars 5µm.

**Figure 5:**
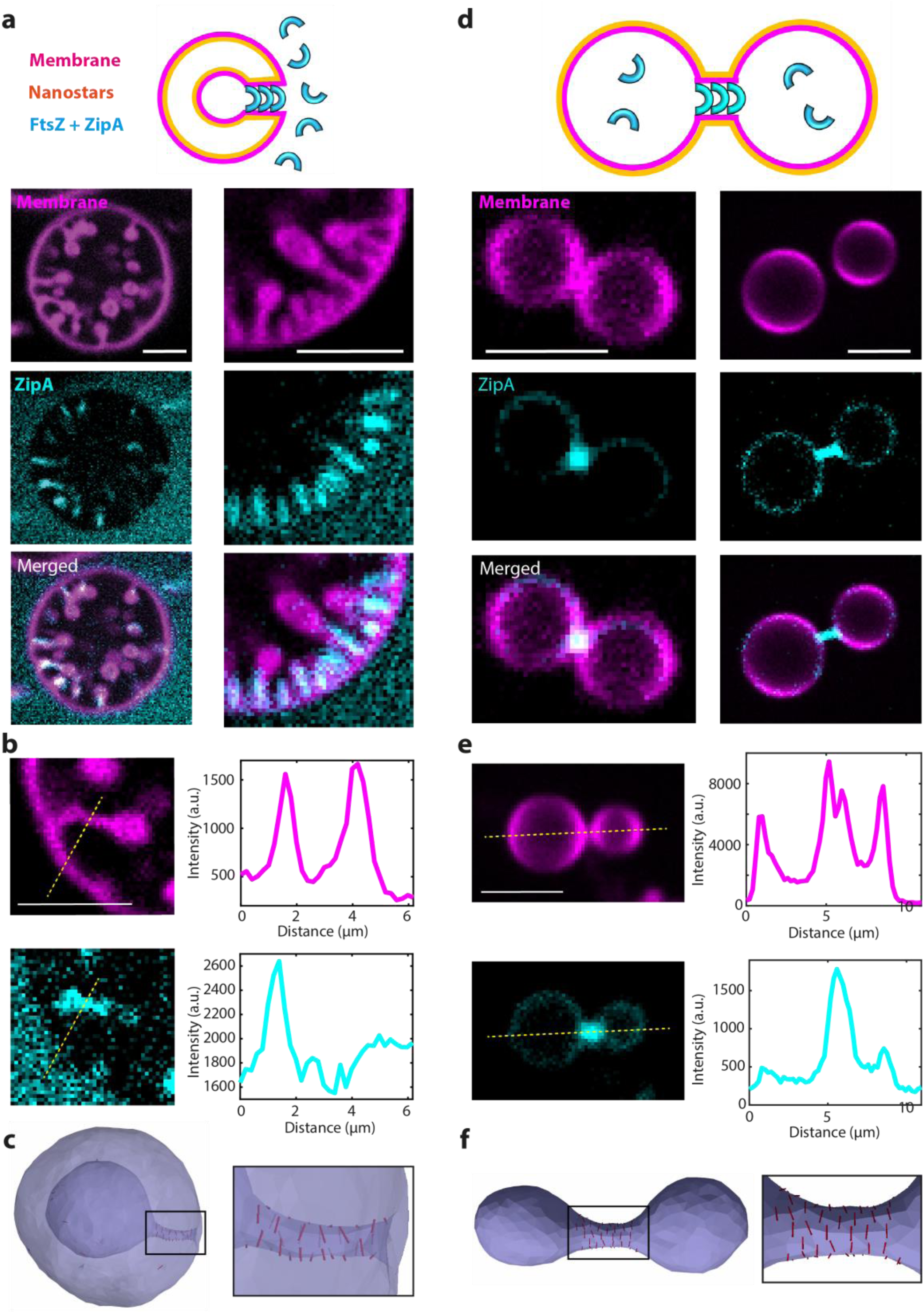
Assembly of FtsZ:ZipA on stomatocyte- and dumbbell-shaped liposomes. **(a)** Assembly of FtsZ:ZipA:GTP on stomatocytes generated with SMS. Schematic on the top row clarifies the topology of the protein and the nanostars with respect to the membrane. Lower panels show representative images. **(b)** Line scan analysis of FtsZ:ZipA enrichment at necks of stomatocytes. **(c)** DTS performed for a fixed reduced volume and 2% protein coverage on the outside membrane show that proteins induce negative curvature. Each red line represents one protein and its orientation on the membrane. Similar behavior was obtained when the volume was reduced by applying osmotic pressure difference, see Figure S9. **(d)** Assembly of FtsZ:ZipA:GTP inside dumbbells generated with SMS. Schematic on the top row clarifies the topology of the protein and the nanostars with respect to the membrane. Lower panels show representative images from 3 independent preparations. **(e)** Line scan analysis of FtsZ:ZipA enrichment at necks of dumbbells. **(f)**: DTS simulation of a vesicle under osmotic pressure, positive global curvature and 5% protein coverage on the inside membrane. Proteins are seen to induce negative curvature. see also Figure S10. All scale bars 5µm.

We found DynA to be highly enriched at membrane necks of both stomatocytes and dumbbells, see Figure 4A and 4D, and Movie 5. This enrichment directly indicates that DynA is able to sense membrane curvature. Protein enrichment on membrane necks and membrane nanotubes can be quantified by the sorting ratio S_R_,^19^ a dimensionless parameter that indicates the fluorescence intensity of the protein at the neck normalized to that of protein bound on the liposome outer surface (see methods). A sorting ratio of S_R_ > 1 indicates enrichment at the neck. For DynA on stomatocytes, we measured a S_R_ of 8.3 ± 4.8 (N=28; error is SD; Figure 4B). On dumbbells, the S_R_ was even higher but it could not be reliably quantified because of the extremely low binding to the liposome surface (Figure 4E). In line with the experiments, membrane deformation by DTS performed in the presence of curved-shape proteins with an affinity for positive membrane curvature resulted in their accumulation on the short catenoid-shaped necks in both stomatocytes and dumbbells (Figure 4C, 4F). Overall, the results indicate that DynA can autonomously localize to membrane necks, which is consistent with *in vivo* observations^18^ that DynA was found to co-localize at the neck of a dividing cell.

We also tested the assembly of FtsZ, the major structural component of the bacterial Z-ring that has been extensively studied before^20^. *In vitro*, FtsZ is known to assemble on negatively curved membranes^21^. Figure 4A and Movie 6 illustrate the assembly of a FtsZ, along with its membrane anchor ZipA,^22^ on stomatocytes that were formed by SMS. Interestingly, rather than a single internal compartment, the FtsZ proteins induce the formation of an extensive array of inward elongated necks that often end with a small spherical internal compartment. The elongated necks indicate that the FtsZ:ZipA complex is able to actively generate negative curvature. This appears to affect the very process of stomatocyte formation, where likely the process of forming one large internal compartment is stalled by the FtsZ as it stabilizes a narrow neck, and subsequently many invaginations occur, leading to an array of necks. The average S_R_ for the FtsZ:ZipA complex was found to be 4.0 ± 1.8 (N=45; error is SD; Figure 4B). In good agreement with these experimental results, our DTS simulations indicated that protein binding to negatively curved membranes would induce neck elongation in stomatocytes (Figure 4C, S9A, S9B) if a sufficiently large directional curvature and an attractive oligomerization potential between proteins were included. When FtsZ was assembled inside dumbbells, elongated necks were also often observed experimentally (Figure 4D, S10, Movie 7), confirming that FtsZ:ZipA complex can generate negative curvature. We also observed this in DTS (Figure 4F). In dumbbells, we measured a higher average S_R_ of 13.2 ± 9.2 (n=30; error is SD; Figure 4E), Taken together, the data indicate that FtsZ:ZipA complexes assemble preferentially on negatively curved membranes as well as are able to actively generate negative membrane curvature.

## Discussion

We presented a novel approach called SMS to induce large-scale membrane deformation in liposomes, which allows to obtain high yields of stomatocyte or dumbbell shapes. Photobleaching experiments confirmed the membrane topology of these structures and allowed to estimate the 30-130 nm size of their defining feature, the catenoid-shaped pore at the neck. Next to developing SMS, we demonstrated that it provides an excellent novel platform to study the interaction of bacterial proteins with curved membranes at the neck region. We found that Dynamin A is able to sense membrane curvature, leading to strong enrichment at the neck of both stomatocytes and dumbbells. As DynA has been suggested to mediate the last step of membrane abscission during bacterial division, encapsulation of DynA in dumbbells by the SMS may serve as a relatively simple experimental setting for reconstituting synthetic cell division by a bottom-up approach. Moreover, we showed that FtsZ:ZipA is able to actively generate negative membrane curvature. Interestingly, this also demonstrates that the membrane structures obtained by the SMS can be further deformed by forces that are exerted by proteins.

Overall, the findings reported here portray the SMS as a promising platform for reconstitution studies of protein binding to membrane necks and in particular on pores that have a catenoid-like shape, which are notoriously difficult to produce with state-of-the-art *in vitro* reconstitution systems. More complex protein complexes that could be reconstituted with this approach include the Nuclear Pore Complex^23^, the ESCRT-III machinery,^24,19^ and the many proteins involved in cell division in eukaryotes, bacteria, and archaea. Currently, the gold standard in characterization proteins interacting with curved membranes is the tube pulling assay, which allows protein reconstitution on positive curvature ^25^ and, by using more complicated procedures, on negative curvature25,26,^28^. However, tube pulling requires highly specialized equipment and is notoriously challenging to perform. The simple and accessible SMS approach developed here is in many aspects complementary to tube pulling (Table S1) and appears to be particularly suited to study protein assembly on membrane regions with negative curvature. Furthermore, the SMS uniquely reproduces the membrane geometry of a toroidal pore, which cannot be achieved by tube pulling, while not requiring any specialized equipment. We anticipate that the SMS will be widely adopted as a general tool for protein and membrane biophysics studies, with high potential to become part of the division machinery for synthetic cells^29^.

## Acknowledgments

We thank Patricia Bassereau, Lennard van Buren, and Alberto Blanch Jover for discussions, Sabrina Meindlhumer for discussions and proposing DynaminA as a particularly fitting candidate for a minimal divisome, Marc Bramkamp for kindly providing the DynaminA plasmid, Eli van der Sluis and Ashmiani van den Berg for discussions and protein purification; Anders Barth for technical help and Jeremie Capoulade for technical support on confocal microscopy. We acknowledge funding support from the BaSyC program of NWO-OCW and from the ERC Advanced Grant 883684. WP is supported by the Novo Nordisk Foundation (Grant Agreement No. NNF18SA0035142).

